# An unbiased, quantitative and versatile method for determining misaligned and lagging chromosome during mitosis

**DOI:** 10.1101/2020.06.25.172478

**Authors:** Luciano Gama Braga, Diogjena Katerina Prifti, Chantal Garand, Pawan Kumar Saini, Sabine Elowe

## Abstract

Accurate chromosome alignment at metaphase facilitates the equal segregation of sister chromatids to each of the nascent daughter cells. Lack of proper metaphase alignment is an indicator of defective chromosome congression and aberrant kinetochore-microtubule attachments which in turn promotes chromosome missegregation and aneuploidy, hallmarks of cancer. Therefore, tools to sensitively and quantitatively measure chromosome alignment at metaphase will facilitate understanding of how changes in the composition and regulation of the microtubule attachment machinery impinge on this process. In this work, we have developed and validated a method based on analytical geometry to measure several indicators of chromosome misalignment. We generated semi-automated and flexible ImageJ2/Fiji pipelines to quantify kinetochore misalignment at metaphase plates as well as lagging chromosomes at anaphase. These tools will ultimately allow sensitive, unbiased, and systematic quantitation of these chromosome segregation defects in cells undergoing mitosis.

## INTRODUCTION

Faithful segregation of chromosomes relies on their accurate alignment at the metaphase plate during mitosis (Kuniyasu et al., 2018). Errors in this process result in chromosome missegregation and aneuploidy, which is defined as an abnormal number of chromosomes for a given cell-type or organism. Aneuploidy drives the development of cancer, at least in part through promoting chromosomal instability (CIN), a state in which the cell displays a high rate of gain and loss of whole chromosomes (Gordon et al., 2012; McGranahan et al., 2012; Soto et al., 2019; Storchova, 2018; Weaver et al., 2007). CIN is a cause of tumor heterogeneity, is associated with a poor patient prognosis and has been linked to drug resistance (McGranahan et al., 2012). To prevent the emergence of aneuploidy and CIN, eukaryotic cells align duplicated sister chromatids at metaphase in a bioriented (also known as amphitelic) manner. The process by which chromosomes are transported to the spindle equator, known as chromosome congression, is thought to promote mitotic fidelity by ensuring that chromosomes enter anaphase in a spatially coordinated manner, thereby preventing random chromosome segregation (Maiato et al., 2017). By aligning at the spindle equator, chromosomes are forced to commence separation and the subsequent poleward movement from the same relative position. Chromosome alignment also increases the likelihood of kinetochore capture by microtubules from both spindle poles, thus favouring a bioriented geometry for chromosome attachments (Cheeseman, 2014; Hinshaw and Harrison, 2018; Joglekar and Kukreja, 2017; Musacchio and Desai, 2017). Finally, chromosome congression promotes correction of erroneous and unstable attachments; proximity to kinase activity at the spindle poles can result in phosphorylations at the kinetochore that weaken the interaction with spindle microtubules (Barisic et al., 2014; Chmatal et al., 2015; Ye et al., 2015). Therefore, chromosome alignment is critical to the fidelity of mitosis and approaches to systematically and rapidly identify the degree of chromosome misalignment are required.

Chromosome alignment and congression can be followed using live-cell imaging or in fixed samples by indirect immunofluorescence. Alignment is typically visually scored in both cases through counting displacement of chromosomes from the equator of the mitotic spindle. The result is often provided as a binary readout of either aligned or misaligned cells. Alternatively, the alignment phenotype is subdivided into multiple poorly defined categories. This approach is prone to bias and may lack reproducibility as it depends on the subjective decision of what constitutes misalignment.

Currently, methods to reliably and reproducibly quantify chromosome misalignment and missegregaton are limited. One approach described by Lampson and Kapoor, measures and normalizes the distance of individual kinetochores from the nearest spindle pole along the pole–pole axis; while this gives a relative distance for individual kinetochores, this methods currently lacks automation (Lampson and Kapoor, 2005). More recently, Fonseca and Stumpff measured average intensity of each pixel column in the fluorophore channel corresponding to kinetochores along the axis of the metaphase plate and along the entire length of the mitotic spindle (Fonseca and Stumpff (2016)). Here, the output is a distribution profile comprising the intensities of kinetochores along the mitotic spindle. From the Gaussian function fit to this distribution, the full width at half maximum (FWHM) is calculated. Each analysed cell provides a single FWHM value that can be used to plot a sigmoidal curve comprising the fraction of the cellular population analyzed by FWHM. In this manner, statistical comparisons between sigmoidal curves reflect the comparisons between different conditions in an experiment. This method gives a global picture of chromosome alignment in the conditions examined but outputs sigmoidal curves deprived of a biological measurement unit, such as the number of misaligned kinetochores.

Here, we developed a method based on analytical geometry to directly quantify misaligned kinetochores at metaphase and lagging chromosomes in anaphase. The method relies on determining the position of the two spindle poles, followed by the establishment of an alignment region based on segmentation of the area between the poles. Kinetochores lying outside a user defined alignment zone are considered misaligned at metaphase, and those lying inside as lagging in anaphase cells. The same approach was adapted to directly measure the distance of individual kinetochores away from a theoretical metaphase plate. We automated the evaluation of chromosome alignment and lagging chromosomes using a series of user-friendly Image J2/Fiji macros (freely available at https://github.com/Elowesab/elowelab) which can be used to rapidly generate quantitative and robust read-outs for these mitotic defects.

## RESULTS

### CELL GEOMETRY VIZUALIZATION

To determine chromosome alignment defects, cells were arrested in metaphase using routine approaches extensively described in the literature (Fonseca and Stumpff, 2016; Kapoor et al., 2006). Briefly, cells were synchronized in prometaphase using drugs that interfere with microtubule dynamics or formation of a bipolar spindle (such as nocodazole, or the KIF5 inhibitor S-Trityl-L-Cysteine, STLC). This was followed by release into media containing the proteasome inhibitor MG132, which prevents anaphase onset due to maintenance of high CDK1 activity and persistent sister-chromatid cohesion. Chromosomes were allowed to align at the metaphase plate before being fixed and stained for kinetochores (using CREST antibodies, or antibodies against CENP-C or another kinetochore protein) and spindle poles (γ-tubulin or a centrosome marker) as a minimum. To ensure that only the population of cells where the spindle and spindle poles are found in the same focal plane were sampled, cells were additionally stained with α-tubulin to facilitate visualisation of the mitotic spindle (**Fig 1A**). Cells with compromised chromosome congression were not expected to properly align chromosomes at metaphase, which was manifest as a disorganized chromosome mass (visualized by DAPI or HOECHST staining) and misaligned kinetochores (marked here by CREST, **Fig 1A**).

**Figure 1:**
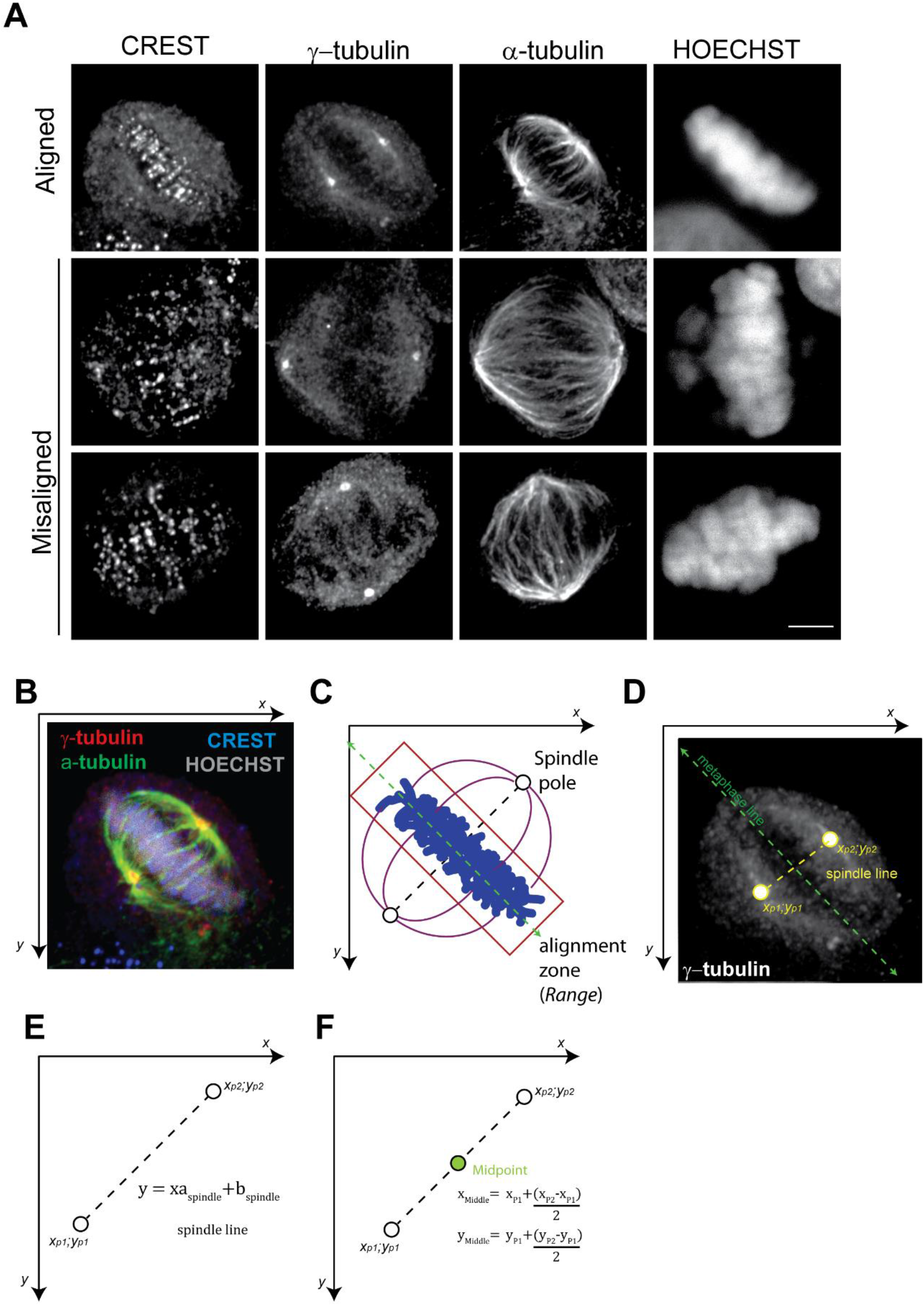
Defining mitotic cells in a Cartesian Plane. A) Representative images of aligned and misaligned chromosomes. HeLa S3 cells were fixed and immunostained with CREST serum to mark the kinetochores, γ-tubulin for the spindle poles, and α-tubulin for the mitotic spindle. The degree of misalignment (second and third row) display a phenotype that is relatively difficult to quantify. Scale bars indicate 5 μm. B) A mitotic cell represented in a Cartesian plane as a function of x and y axes. C) A cartoon representation of the key inputs that are provided by the user: positions of the spindle poles and the parameters for identifying the alignment zone (*Range*, red rectangle) around the metaphase plate (dashed green line). D) Inferring the metaphase line from the position of the spindle poles. The metaphase line is assumed to be perpendicular to the spindle line that passes through the two spindle poles. E) The equation of the spindle line is deduced from the coordinates of the spindle poles in the Cartesian plane. Spindle pole centroids are indicated by white circles. F) The midpoint coordinate (green circle) is deduced from the equation of the spindle line.

### MODELLING CELL GEOMETRY

To facilitate modelling of the geometry, cells are spatially translated into a Cartesian plane (Fig. **1B**). This allowed mathematical definition of an alignment zone encompassing the theoretical metaphase plate with kinetochores localized outside this region considered unaligned (Fig. **1C**). To achieve this, we developed an approach that is broadly divided into three steps: identification of spindle pole position, determination of a user defined alignment zone, and finally, enumeration of kinetochores outside the defined alignment zone.

After the cell of interest is selected, the method relies on two main inputs from the user, required at the beginning of the operation. The first is to identify the position of the two spindle poles in the image (**Fig. 1D)**. The significance of the poles lies in the assumption that a theoretical metaphase plate (represented by the metaphase line, green in **Fig. 1D**) is perpendicular to and intersects at the midpoint of the line connecting the two spindle poles (the spindle line, yellow in **Fig. 1D**). To this end, *(x*_*P1*_*; y*_*P1*_) and *(x*_*P2*_*; y*_*P2*_) represent the coordinates of the first and second spindle poles, respectively. Given the two points, the equation of the spindle line that passes through both poles and thus the midpoint in between the poles can be deduced (**Fig. 1 E, F**). This midpoint represents the centroid of the alignment zone.

The second user-defined input is a pair of values that are used to determine a variable we call the *Range*. For identifying chromosome alignments defects, the *Range* essentially corresponds to the alignment region, which is eventually excluded in order to quantify misaligned kinetochores lying outside this zone. Based on the premise that metaphase is a polygon centred around the spindle midpoint (**Fig 2A**), the user inputs 2 *Range* parameters which we call the “total segments” and “aligned segments” to define the width of the *Range* relative to the distance between the poles. Here, the value for “total segments” represents the number of equal segments into which the area between the spindle poles is divided, whereas “aligned segments” is the number of central segments encompassing the desired *Range*. In this manner, the line between the poles is divided into equal segments, where the segment(s) in the middle represent the *Range* (**Fig 2B**). The values for both *Range* parameters are arbitrary and user-defined, with some limitation. To ensure that the alignment zone is central, if the spindle length is divided into an odd number of segments, the alignment zone should also be divided into an odd number of central segments. In contrast, if the total number of segments is an even number, the value for aligned segments should also be even. For instance, if the spindle length is divided into five equal segments, the narrowest *Range* can be represented by the single central segment. A broader metaphase may be required in which case the three central segments would be considered as the alignment zone, and the value provided for “aligned segments” would be 3.

**Figure 2:**
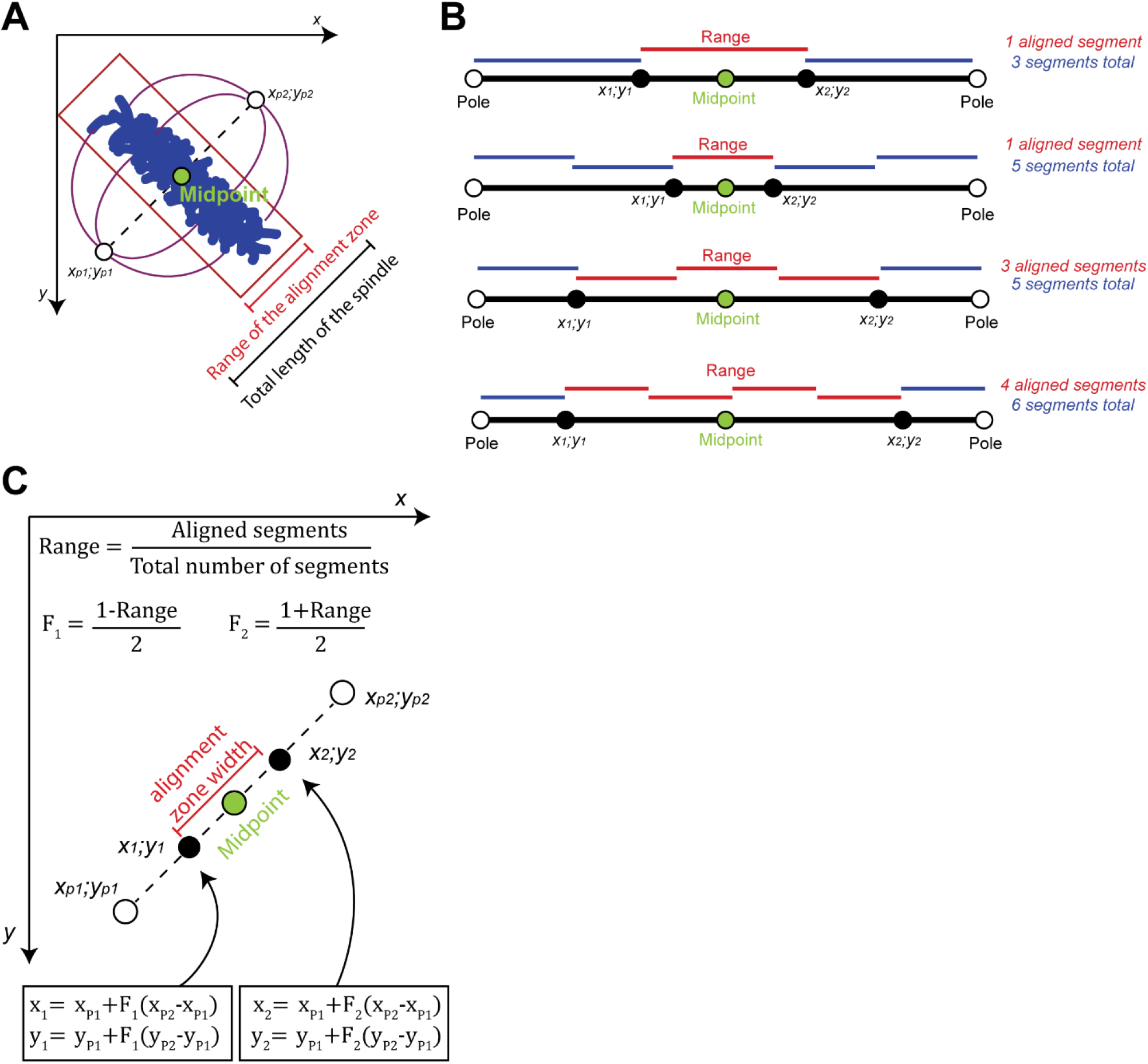
Identifying chromosome alignment regions as a function of the *Range*. A) Scheme depicting the *Range* and the theoretical alignment region as a function of the pole-pole distance. B) Examples illustrating variations of the *Range* segmetation scheme. The poles are represented by white circles, the midpoint by the green circles, and the points delimiting the alignment region are represented by black circles. The poles are used to delimit a line that is divided by an arbitrary number, creating several sections. One or more sections are chosen as the aligned sections. C) Graphical representation of the *Range* relative to the spindle poles and the midpoint. The equations for the calculation of the points (*x*_1_*;y*_*1*_) and (*x*_2_*;y*_*2*_) delimiting the alignment region (the black circles) are shown.

Likewise, if the spindle line is divided into six equal segments, where the four central sections are considered by the user to be the alignment region, then the values for “total segments” is 6, and 4 for “aligned segments” (**Fig. 2B**). The choice for the number of segments in each category would thus allow for precise definition and calibration of the *Range*. In this manner a graded or subtle alignment defect can be reliably identified. After defining pole coordinates and setting the *Range* parameters, the points that delimit the alignment region along the spindle line (ie. the width of the alignment zone); defined as *(x*_*1*_*; y*_*1*_) and *(x*_*2*_*; y*_*2*_) in the Cartesian plane, are calculated as a function of the *Range* based on analytical geometry (**Fig. 2C**, see methods).

### DEFINING THE ALIGNMENT ZONE POLYGON IN THE CARTESIAN PLANE

Once the points that delimit the alignment zone width along spindle line are determined, the outer limits of this region are defined by establishing the points that delimit the rectangular alignment zone centred at the midpoint (**Fig. 3A**). To do this, lines parallel to the metaphase line and that pass though *(x*_*1*_*; y*_*1*_) and *(x*_*2*_*; y*_*2*_), defined here as alignment lines, are calculated (**Fig 3B, C**). The width (W) of the rectangle is set by the position of the alignment lines, but its length (L) is not a function of the spindle and here, we set them to incorporate the entire cell as metaphase plate dimensions can vary (**Fig. 3D**). By applying the endpoints described above to the equations of the alignment lines, the four points of the rectangle defining the alignment zone are obtained (see methods). Because the slope of the spindle line in the Cartesian plane can be positive or negative, the endpoint values on the X and Y axis of the plane used to determine the limits of the rectangle are set accordingly in the macro described below (**Fig 3 D, E**). In sum, our geometrical model of the cell requires as minimal inputs by the user the coordinates of the spindle poles if they are being manually defined, and arbitrary *Range* parameters to determine the alignment region.

**Figure 3:**
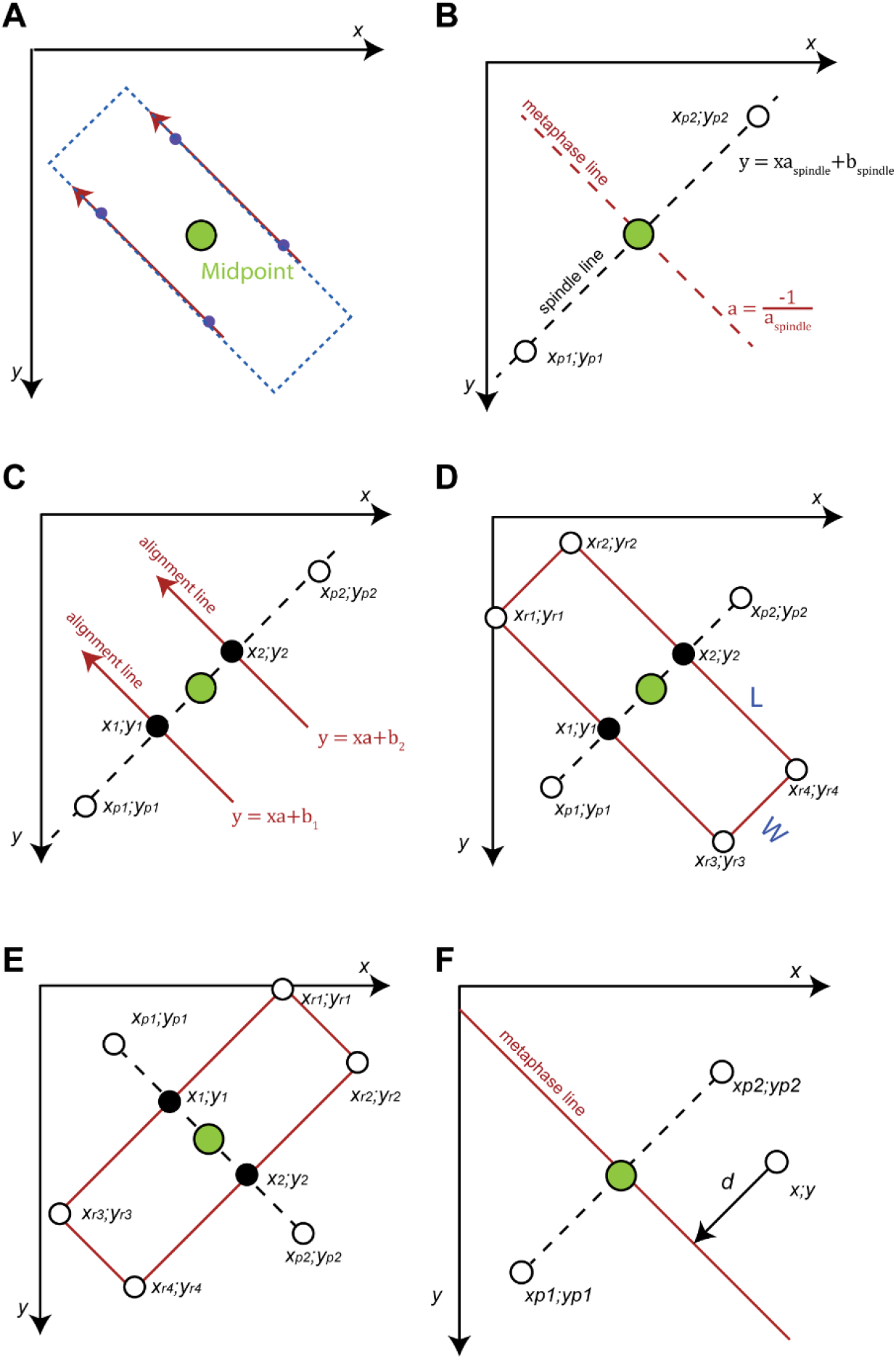
Geometrical model of the chromosome alignment region in a Cartesian plane. A) The alignment region is defined a polygon centred around the midpoint. B) The metaphase line is perpendicular to the spindle line. The slope of the metaphase plate is deduced from the spindle line. C) Two points (black circles) along the spindle line delimit the width of the alignment region. The red alignment lines passing through these points delimit the alignment region and are parallel to the metaphase line. As a result, both alignment lines and metaphase line have equal slope coefficients. D) To define a polygon encompassing the alignment zone, the width is limited by the alignment lines, and the extremities of the image are used to limit the length, ensuring that the whole cell is encompassed. In this configuration, a<0 and the first corner coordinate (*x*_*r1*_*; y*_*r1*_) is defined as the point where the first alignment line crosses the y axis. The second point (*x*_*r2*_*; y*_*r2*_) is the closest point on the second alignment line to (*x*_*r1*_*; y*_*r1*_). For the third point (*x*_*r3*_*; y*_*r3*_) a value greater than the image size was considered, and the closest point to it in the second alignment line defines the fourth point (*x*_*r4*_*; y*_*r4*_). E) If the metaphase plate has negative slope a>0, the first coordinate is calculated by the point where the first alignment line intercepts the x axis. A similar rationale to the previous configuration is used to find all points delimiting the rectangle. F) Since each kinetochore is separately counted in Fiji, their individual position can be determined by built-in functions. The distance of each kinetochore from the theoretical central line in between the poles (metaphase line) can then be calculated.

The method described above classifies kinetochores as either aligned or misaligned. However, the distance between kinetochores and the metaphase line can give an alternative and independent readout for chromosome alignment (**Fig. 3F**). To measure kinetochore distances, the metaphase line is defined as indicated above, based on the position of the spindle poles. The kinetochores are then either manually or automatically detected and the shortest distance between individual kinetochores and the metaphase line is computed (see methods).

### SEMI-AUTOMATED IMAGE J MACRO FOR QUANTIFICATION OF MISALIGNED CHROMOSOMS

In order to implement and automate the above methods for detection of kinetochore misalignment, we created Fiji (ImageJ2) macros to count kinetochore misalignment as well as measure distances of individual kinetochores away from the metaphase line. The first step of the macro for kinetochore misalignment is a prompt for the user to open images (single plane or a projected stack) that will be used to identify the spindle poles, kinetochores and the chromatin in that particular order which is predetermined to facilitate automation of the method. Next, the user is prompted to select the region/cell of interest (**Fig. 4Ai**). Signals outside of this region are subsequently excluded to ensure that only kinetochores from the cell of interest contribute to the final kinetochore count. Once the region of interest is selected, the user is prompted to set the parameters necessary to determine the *Range*; the number of “total segments” and “aligned segments”. The y-axis limit which defines the length of the alignment zone, is set to a default 15000 pixels, and can be manually changed if desired, depending on the parameters used for image acquisition (**Fig. 4Aii)**. The next step is to identify the position of the spindle poles. The user is prompted to identify the poles either automatically or manually in the appropriate channel (**Fig. 4Bi**). Automatic detection is accomplished with the built-in IMAGE J function *Analyze Particles*, as indicated in the methods (**Fig. 4Bii**), whereas the multi-point function is used for manual selection. The position of the poles as well as *Range* parameters are then used to determine the alignment zone (**Fig. 4Ci**), the signals inside this zone are deleted with the *Clear* function in the kinetochore image, and only the objects (misaligned kinetochores) outside the alignment region remain (**Fig. 4Cii**). These kinetochores are then counted automatically with ImageJ/Fiji built-in functions. The user is prompted to evaluate and can manually threshold the remaining signal in the kinetochore channel to facilitate accurate quantification of the number of objects remaining (**Fig. 4Ciii-iV**).

**Figure 4:**
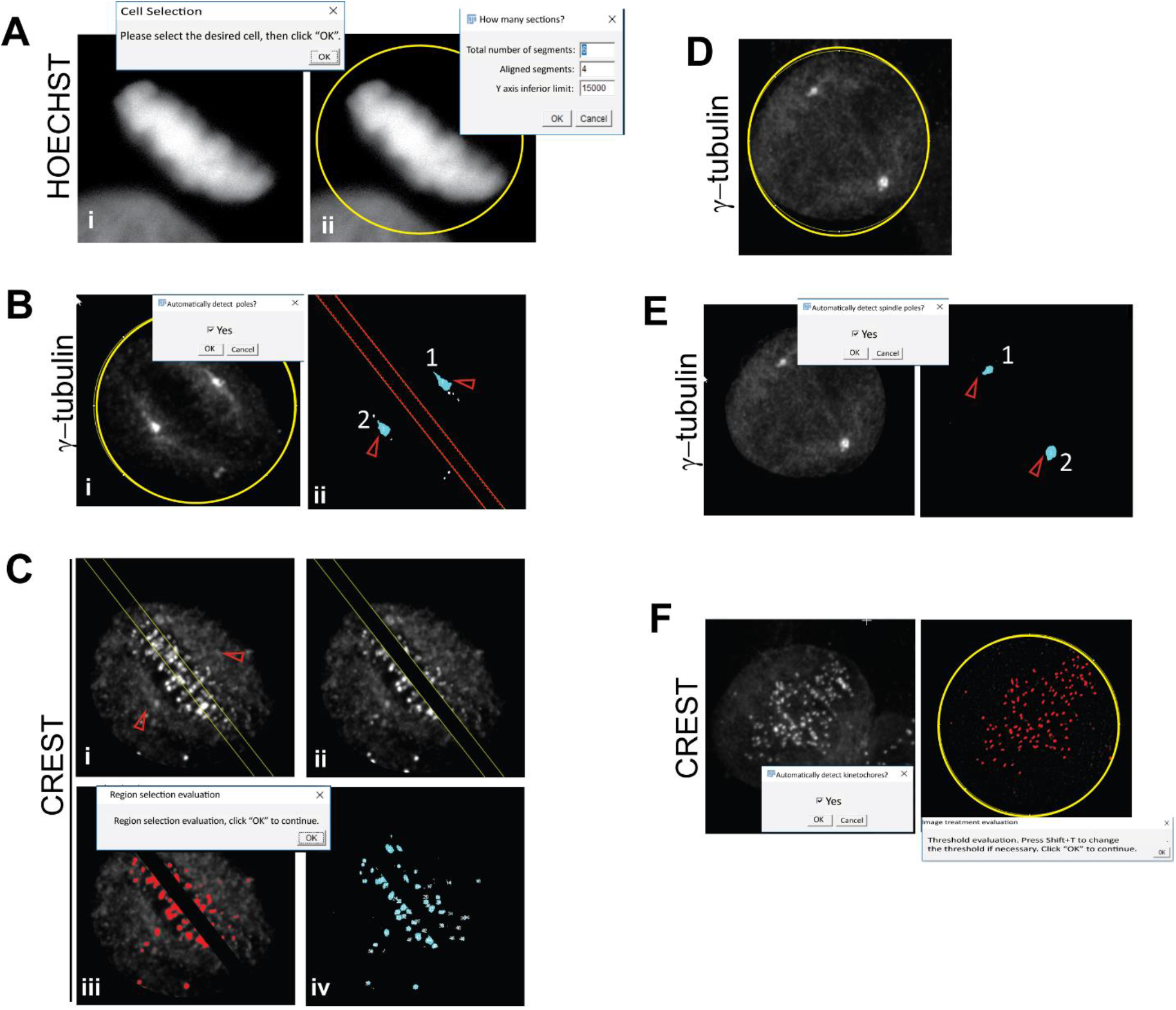
Key steps in the execution of the chromosomes alignment and kinetochore-metaphase distance macros. A) (i) The initial step is selection of the ROI in the DAPI/Hoechst channel. Signal outside this selection is deleted from the CREST and γ-Tubulin images. (ii) This is followed by selection of the *Range* parameters. B) (i) Selection of the spindle poles is then initiated in the appropriate channel (here, in the image stained for γ-tubulin signal. (ii) The spindle poles (red triangles) are then detected automatically in Fiji by the built-in function *Analyze Particles* (right panel). The centroid from each area is used as pole coordinates for the mathematical operations. C) (i) The alignment region is delimited by the dotted lines in the image corresponding to kinetochores. (ii) In the chromosome alignment macro, the signal inside the selection is removed, with only the signal from misaligned kinetochores remaining. (iii) The *Threshold* function is then used to identify the remaining kinetochores. (iv) These are then counted in the macro using *Analyze Particles* functions. D) To measure kinetochore distance from metaphase, ROI selection occurs in the image corresponding to the spindle poles. As in A) signals outside the ROI are deleted in the images corresponding to the poles and kinetochores. E) Spindle poles are detected as in B, F) All kinetochores (after thresholding) are detected and their distance from the metaphase line is computed.

Measurement of kinetochore distances does not require definition of an alignment zone or *Range* parameters. Instead, after the poles are identified (either automatically or manually) and the metaphase line computed as above (**Fig 4D, E**), the user is prompted to choose between manual or automatic kinetochores detection (**Fig. 4 F)**. Once identified, the distance of every kinetochore to the metaphase line is then computed (see methods).

### EVALUATION OF CHROMOSOME ALIGNMENT IN AURORA B AND CENP-E INHIBITED CELLS

To generate chromosome misalignments and reliably test our pipeline, we took advantage of the phenotypes associated with the inhibition of two key proteins that regulate chromosome alignment during mitosis: the kinase Aurora B and the kinesin CENP-E. Aurora B (the catalytic subunit of the chromosomal passenger complex) localizes mainly to the inner centromere of chromosomes during early mitosis where it phosphorylates multiple targets at kinetochore proximal centromeres and inner kinetochores to ultimately regulate the microtubule-kinetochore interaction status (Lampson and Grishchuk, 2017; Welburn et al., 2010) (Hindriksen et al., 2017). Moreover, Aurora B activity can destabilize attachments by inhibiting protein phosphatases responsible for stabilizing microtubule binding (Foley et al., 2011; Liu et al., 2010; Meppelink et al., 2015; Nijenhuis et al., 2014). CENPE is a processive motor protein that moves its chromosome cargo along spindle microtubules in a plus-end directed fashion thereby permitting congression to metaphase (Espeut and Abrieu, 2015; Schaar et al., 1997; Yu et al., 2019). CENP-E is also thought to promote the formation of end-on stable kinetochore-microtubule attachments to maintain chromosome alignment at the metaphase plate (Shrestha and Draviam, 2013). Inhibition or depletion of either Aurora B or CENP-E in human cells causes alignment defects and lagging chromosomes in anaphase (Bennett et al., 2015; Cimini et al., 2006; Ditchfield et al., 2003; Harborth et al., 2001; Hauf et al., 2003; Tanudji et al., 2004; Yen et al., 1991).

To evaluate the accuracy and robustness of the analysis pipeline, we used it to determine chromosome alignment and metaphase kinetochore distances in cells where Aurora B or CENP-E were inhibited with ZM447439 or GSK923295, respectively, using the lowest working concentrations commonly found in the literature (See **Fig. 5A** for the experimental timeline). HeLa-S3 cells were used for these experiments as this is a standard cell line routinely utilized for studies of mitosis. Measurement of the distances of individual kinetochores from the metaphase line confirmed a significant increase in kinetochore distances from the metaphase line of cells treated with either 0.5 μM ZM447439 or 10nM GSK923295 as expected (**Fig. 5B,D**). Similar conclusions were reached when the mean kinetochore distance was calculated for each cell examined (**Fig. 5C**).

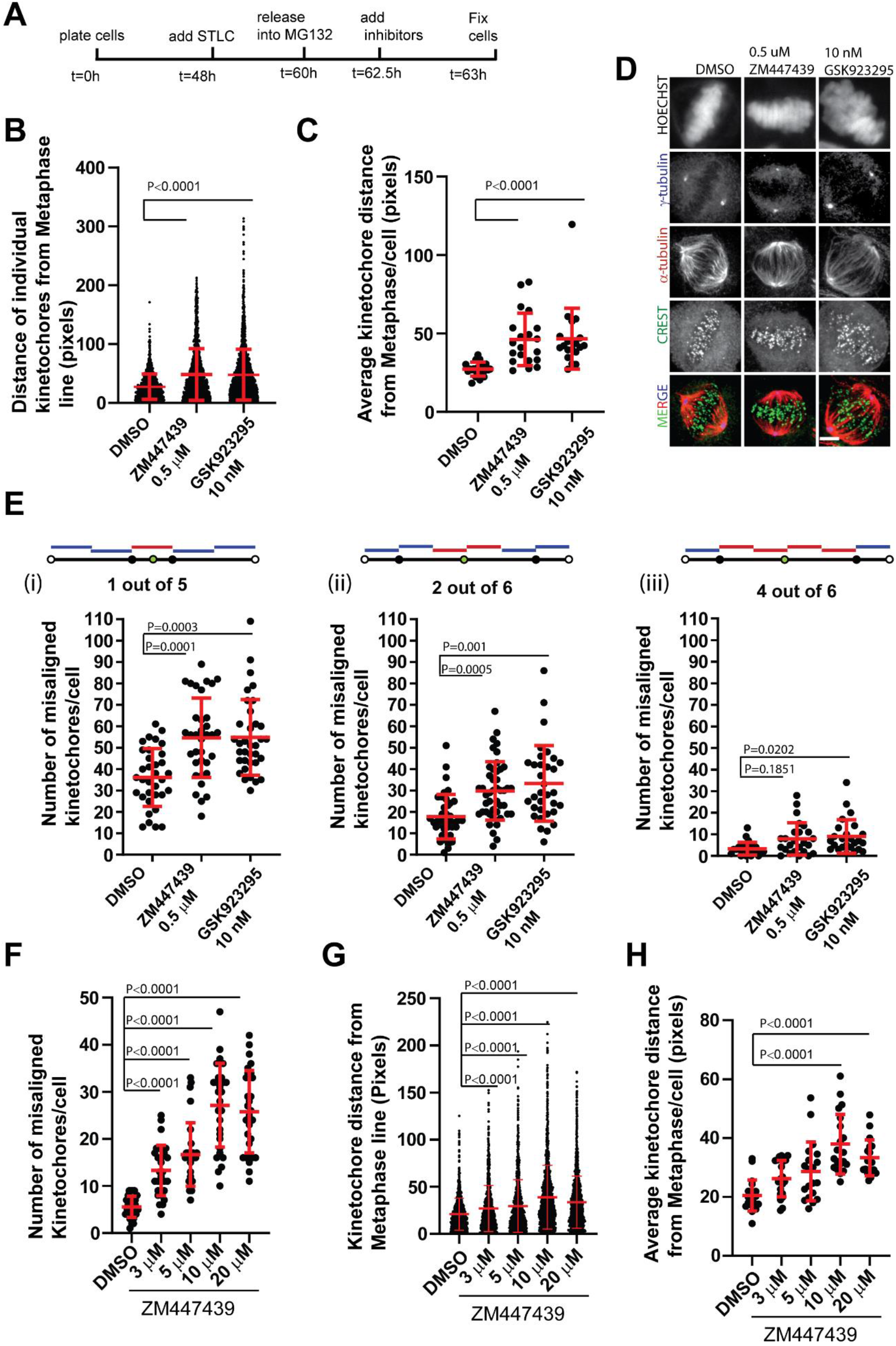
Validation of the chromosome alignment and kinetochore-metaphase distance operations and macros. A) Experimental timeline for validation of the chromosome alignment kinetochore to metaphase distance pipelines. B) Distance of individual kinetochores from the metaphase in cells treated as indicated. Data is from a representative experiment (N=3), with a minimum of 20 cells per condition. C) Average kinetochore distance per cell for the Data in B. D) Representative images of the data shown in B, C. E) Measurement of kinetochore misalignment under the conditions indicated using *Range* parameters of (i)1 ou of 5, (ii) 2 out of 6, and (iii) 4 out of 6. F) The number of misaligned kinetochores, (G) kinetochore distance from metaphase, and (H) Average kinetochore distance from metaphase in cells treated with increasing doses of the Aurora B inhibiter ZM447439. A range of 4 out of 6 was used for evaluation of the alignment errors. Statistical significance is tested by non-parametrical ANOVA (Kruskal-Wallis test), and conditions were compared through Dunn’s multiple comparison test.

To determine kinetochore misalignment 3 variations of the *Range* were used, each with progressively increasing width (**Fig. 5E**): (i) 1 aligned segment out of 5, (ii) 2 aligned segment out of 6, and (iii) 4 aligned segments out of 6. As expected, low levels of misaligned chromosomes were observed in cells treated with DMSO, while cells treated with either inhibitor presented a marked and statistically significant increase in chromosome misalignment (**Fig. 5E**) for almost all *Range* values (see below). In addition, the absolute number of misaligned kinetochores counted from the same dataset decreased with increasing alignment zone (*Range*) size for all conditions tested (compare the values for the same condition between Fig **5Di, ii** and **iii**). This is as expected, given that a wider alignment zone/*Range*, would result in more kinetochores being considered as aligned, further validating our approach. Moreover the data also confirm that the choice of values for the *total segments* and *alignment segments* inputted determines the extent of misalignment and should be optimized by the user. For example, using a *Range* of 1 out 5 and 2 out of 6 revealed differences in chromosome misalignment with higher statistical significance compared to a *Range* of 4 out of 6 in cells treated with either ZM447439 or GSK923295.

To provide further support for the robustness of the method in assessing diverse degrees of chromosome alignment, we measured the kinetochore distances from metaphase and quantified chromosome misalignment in cells treated with increasing concentrations of ZM447439. Using the macro and selecting the arbitrary *Range* ratio of 4 aligned segments out of 6, the average number of misaligned kinetochores increased incrementally with increasing doses of ZM447439 until a plateau was reached at 10uM (**Fig. 5E**). In agreement, measurement of kinetochore distances also demonstrated larger displacement from the metaphase line with increasing doses of the Aurora B inhibitor, at the level of individual kinetochores as well as individual cells (**Fig. 5F, G**). Collectively, these observations strongly correlate with the effect of these inhibitors on chromosome alignment documented in the literature and validate the methodology presented above.

### EVALUATION OF LAGGING CHROMOSOMES

The translation of the cell into a Cartesian plane and its segmentation can be simply and rapidly adapted (and automated in Fiji/Image J2) to measure lagging chromosomes in anaphase. In this context, instead of using the poles to delimit the spindle line, the two anaphase chromosome masses are used (**Fig 6A**). After selecting the region of interest, the user is again prompted to define the *Range* as above. Next, the center of each chromosome mass in the DAPI/HOECHST image manually or automatically is detected (see methods). Once these anchor points have been defined, the same geometric procedure described above is used to delimit the user-defined *Range* in the center, which now represents the region of interest for evaluating the number of lagging chromosomes and is thus retained, whereas the area outside is excluded To validate this method and the associated macro, we treated HeLa-S3 cells entering mitosis with ZM447439 to generate chromosome attachment errors (**Fig 6B**). Quantification of anaphase lagging chromosomes in control and ZM447439-treated cells using the macro and the user-defined segmentation of the area between the divided chromosome masses (in this case, a *Range* of 3 out of a total of 5) confirmed these observations and demonstrated a statistically significant difference in the number of lagging chromosomes between control and ZM447439-treated cells (**Fig 6C,D**).

**Figure 6:**
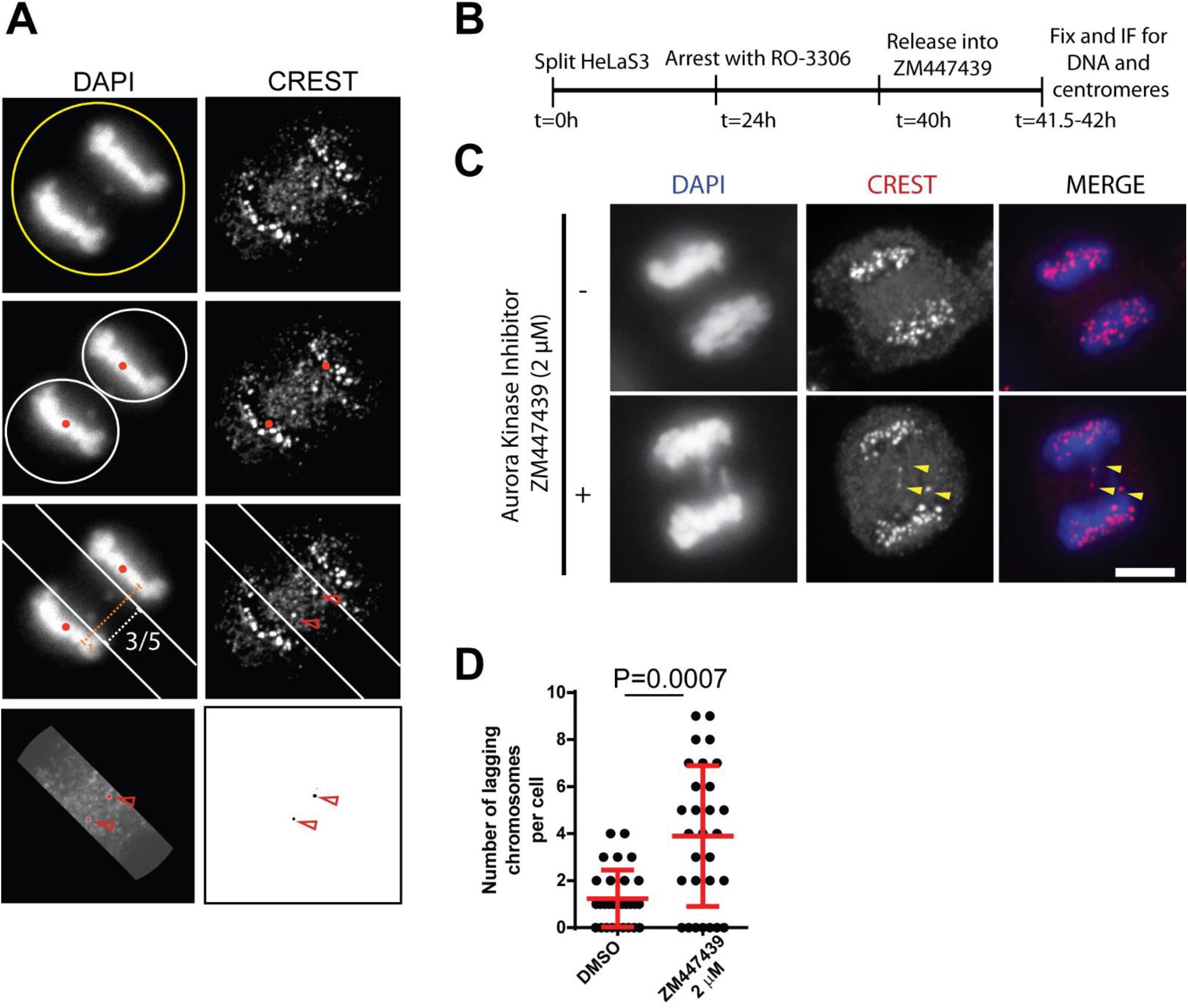
Adaptation and of the Range operations for the measurement of lagging chromosome. A) Steps used by the macro to count the number of lagging chromosomes. After the desired cell is selected and outside signal deleted, the center of each anaphase mass in the chromatin channel is determined using Fiji built-functions. The lagging region is determined as a function of the *Range*, and signals outside this are excluded in the image corresponding to the kinetochores. Kinetochore thresholding and detection then facilitate counts of objects/kinetochores within the zone defined by the *Range*. B) Experimental timeline for validation of the lagging chromosome macro. C) Representative immunofluorescence images from the experiment outlines in B. Kinetochores from lagging chromosomes are marked with yellow arrowheads. Scale bars indicate 5 μm. D) The number of lagging chromosomes per cell in cells treated as indicated. The experiment was assessed with a *Range* of 3 out 5, which is represented in A. Statistical significance is tested by non-parametrical t-test (Mann Whitney test).

## DISCUSSION

Here, we used a Cartesian plane model of the cell and analytical geometry to facilitate the detection and quantification of misaligned kinetochores and anaphase lagging chromosomes, phenotypes commonly associated with mitotic defects and often scored in the literature. Moreover, we have automated the operations in a series of user friendly, flexible macros (https://github.com/Elowesab/elowelab). We have validated this approach and the associated macros using data acquired from cells treated with Aurora B and CENP-E inhibitors, both of which have been consistently reported to induce perturbations in chromosome alignment at metaphase and their segregation at anaphase (Cimini et al., 2006; Ditchfield et al., 2003; Tanudji et al., 2004; Yen et al., 1991). Our results are fully consistent with the literature and confirm the utility of the approach developed here for automated and unbiased scoring of these defects.

In addition to providing an objective quantification of chromosome alignment errors, one of the key strengths of this approach is the versatility in *Range* parameters which allows for standardized definition of the alignment zone. *Range* parameters can be tailored and optimized by the user to help improve inter-experiment variation, for example. Moreover, considering that the *Range* is a constant fraction of spindle length, the inherent variability of different cell sizes in a heterogeneous cell population is accounted for in the execution of the method. These characteristics will undoubtedly reduce user time and effort spent on image processing post-acquisition as well as data normalization.

We show here also that the method is flexible and can be adapted to measure lagging chromosome in anaphase (Fig. 6). Furthermore, although limited to images of fixed cells in this manuscript, this approach can in principle be expanded to analysis of videomicroscopy of cell division; by determining the poles and kinetochore positioning in each frame from a live-cell movie, the dynamics of kinetochore positioning in relation to a theoretical metaphase plate can also be inferred. If the relative position of each kinetochore is calculated for each time point it would be possible to impartially evaluate chromosome congression in real time.

One limitation of the approach presented here is that it assumes a symmetric spindle with the alignment region centred between the two half-spindles and thus misalignment from malformed or asymmetric spindles may not be accurately detected. Additionally, as with any other automated or semi-automated image analysis methods, reliable immunostaining (of kinetochores and spindle poles in this case) are key to generating reproducible results. However, this is countered by the flexibility offered in detection of the anchor points (spindle poles and the anaphase DNA centroids), which in the macro can be identified either manually or automatically with built-in Fiji functions. This feature may be useful in the absence of pole staining, or in cases where poor staining of centrosome or spindle pole markers in the image makes automatic detection problematic. In this case, pole coordinates could be inferred (albeit less precisely) from mitotic spindle architecture and positioning using α-tubulin staining, for example. Moreover, manual selection of kinetochores and threshholding at key points during pauses in the operation allow for further user optimization, as necessary. In conclusion the method outlined in this study provides a reliable, unbiased, semi-automated and versatile tool for assessment of the most common mitotic phenotypes and will serve as a useful tool for researchers in the cell cycle and aneuploidy fields.

## MATERIAL AND METHODS

### CELL CULTURE AND IMMUNOFLUORESCENCE

HeLa-S3 cells were cultured in DMEM (Hyclone) supplemented with 10% (vol/vol) of bovine growth serum and Pen/Strep (100 μg/ml, Hyclone) at 37°C and 5% CO_2_. For measuring chromosome alignment, cells seeded onto coverslips were arrested in prometaphase with STLC (Sigma, 5 μM) for 12 h. Subsequently, cells were released into MG132 (Calbiochem, 20 μM) for 2.5 h then fixed with PTEMF (0.2% Triton X-100, 20 mM PIPES pH 6.9, 1 mM MgCl_2_, 10 mM EGTA and 4% formaldehyde). Prior to fixation, cells were treated with either Aurora B (ZM-447439, Enzo) or Cenp-E (GSK923295, Selleckchem, S7090) inhibitors at the indicated concentrations for 30 minutes, before being fixe and stained. For measure lagging chromosomes in anaphase, cells were arrested or 16h in G2/M with 4μM RO-3306 before being released for 1.5-2h into control media or media containing ZM447439. Primary antibodies for immunofluorescence were used at 1 μg/mL, as follows: CREST anti-Centromere serum (HCT-0100, Immunovision), anti-α-Tubulin (DM1A, Santa Cruz) and anti-γ-Tubulin (T3559, Sigma-Aldrich). HOECHEST 33342 (Thermo Scientific) was used at 1 mg/mL.

### MICROSCOPY

Image acquisition was performed with a with an Olympus IX80 on an inverted confocal microscope equipped with a WaveFX-Borealin-SC Yokagawa spinning disc (Quorum Technologies) and an Orca Flash4.0 camera (Hamamatsu). Metamorph software (Molecular Devices) was used to acquire images 16 z-stacks at 0.2 μM intervals that were then projected onto a single plane for further processing. Optical sections were acquired with identical exposure times for each channel within an experiment.

### MACRO AND IMAGE ANALYSIS

Image analysis was performed using ImageJ2 (Fiji version 1.52i). The initial steps integrated into the macros provided with this manuscript are as follows: First, upon starting the macro, the user is prompted to open the single plane or projected image files of the kinetochore, chromosomes and/spindle poles in a particular order which is retained to facilitate automation. Next, the cell of interest is selected using the *Oval* tool, followed by the deletion of signals outside the cell with the *Edit>Clear outside* command. The user is then prompted to input the *Range* parameters: Total number of segments and aligned segments as well as the Y-axis inferior limit. For quantifying the number of misaligned chromosome (chromosome alignment.ijm), the selection of each pole is prompted by the macro, either manually or automatically. If the user choses to manually select the poles, the *Multi-point* tool (right click on the point tool to select multi point) is used. The macro then initiates the *Analyze>Measure* command to output the centroid coordinate for each pole. Alternatively, if the user chooses the option of automatic detection of the poles, the macro initiates the *Analyze>Analyze Particles* command. Once the poles and *Range* are defined, the four points defining the corners of the alignment region are calculated (see below) and the resulting polygonal selection created in the image corresponding to the kinetochores. Signal from inside the polygon are subsequently deleted with the *Clear* function resulting in an image containing only the misaligned kinetochores outside the alignment zone. The kinetochore image is filtered using the *set Autothreshold* function, followed by *Make Binary*, and *Watershed*. Further manual thresholding at the discretion of the user can be implemented to optimize kinetochore detection. The number of kinetochores/objects and their positions are detected with the *Analyze Particles* function. As with thresholding, the parameters for the *Analyze Particles* function can be additionally optimized by the user. The final output from the macro is a summary table with the number of misaligned kinetochores as well as their average size, and a second table with a detailed description of for each kinetochore identified including the area and position.

For measuring kinetochore distances away from metaphase (kinetochore to metaphase distance.ijm), the selection of each pole is performed as described in the previous paragraph following the parameters inputted by the user for the *Range*. The image is then filtered through *Process>Filters>Convolve* and *Image>Adjust>Threshold>MaxEntropy dark*, followed by the detection of kinetochores with the *Analyze Particles* function. The shortest distances of individual kinetochores from the metaphase line are calculated (see below), and the final output are two windows: the first containing the kinetochore identification number and its distance to the metaphase plate. The second table lists the distance between the poles.

Finally, in the macro to identify lagging chromosomes in anaphase cells (lagging chromosomes.ijm), the desired cell is selected and the exterior is deleted as indicated above. The coordinates at the centeroids of the two anaphase chromosome masses can be selected manually using the *Multi-point tool* as indicated above in the image of the chromatin signal. Alternatively, the center of the anaphase chromosome masses is determined automatically by selecting the first mass, excluding the exterior image with *Clear*, and then using the *Threshold* function and *Create selection*, to create a selection containing the first anaphasic chromatin mass. The *Measure* function will then output the centroid coordinates corresponding to the center of the chromatin mass. The same procedure is repeated in the macro for the second chromatin mass. After obtaining both coordinates and providing *Range* parameters, the polygon can be calculated and in the kinetochore image with the built-in *Polygon* function. The function *Clear outside* is then used to exclude all signal apart from the kinetochores inside the “lagging region” defined by the polygon. The kinetochores are counted with the *Analyze particles* function as indicated above in for measuring chromosome alignment. The output includes table with the number of lagging chromosomes, total area and average size, and a second table that list the area and position of the chromosome masses.

### MATHEMATICAL DEDUCTIONS

To determine the coordinates of the spindle, the cell is interpreted in a Cartesian plane. Given the two poles where (*x*_*P1*_, *y*_*P1*_) and (*x*_*P2*_, *y*_*P2*_) represent the coordinates of the first and second spindle poles, respectively, the equation of the line that passes through the poles (ie. spindle line) is represented by:

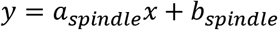

Where the slope of the spindle line (*a*_*spindle*_) is given by the equation:

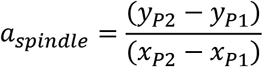

Using this equation, *a*_*spindle*_ is calculated, and is then used to determine the slope of the metaphase line (*a*) which is perpendicular:

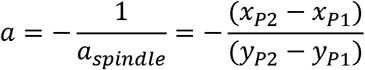

To define the alignment zone, the *Range* parameters are user-defined as follows:

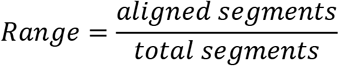

Where ‘aligned segments’ indicates the number of central segments that constitute the alignment zone and ‘total segments’ is the total number of segments that the area between the spindle poles is divided into. The offset factor (F, see Fig. 2C) defines the points along the spindle line that demark the boundaries of the alignment zone and is a function of the Range. Thus:

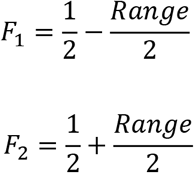

These functions are then used to determine the points on the spindle line through which the alignment lines pass, as follows:

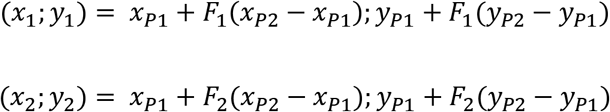

With the deduction of the two points delimiting the alignment region and the slope of the metaphase line, the equation of the alignment lines can be determined as in Fig 3C. Given that these lines are parallel to the metaphase line, their slope can be represented by *a.* Thus, the two alignment lines that determine the width of the alignment region are define by the following equations:

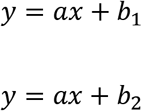

Where:

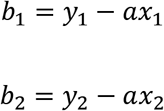

Next, the four points on the alignment lines that limit the outer corners of alignment polygon are deduced (**Fig 3D,E**). If a>0, then *x*_*r1*_*;y*_*r1*_ is set to *0;b*_*1*_. *x*_*r1*_ set at 0 signifies the limit of the image in the Cartesian plane, where it intercepts the y-axis **(Fig. 3D**).

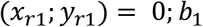

The second coordinate (*x*_*r2*_*; y*_*r2*_) is the closest point on the parallel alignment line. This is calculated by:

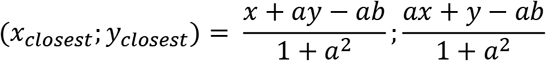

By applying the above equation, *x*_*r2*_*; y*_*r2*_ is defined:

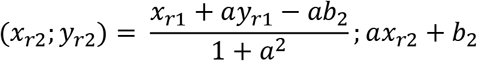

The third point is defined by the upper limit of the y axis in the Cartesian plane. In the macro, this is set to 15000 pixels by default. Thus, the third point is calculated as:

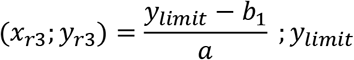

The fourth and final coordinate can be calculated by determining the closest point on the second line to *x*_*r3*_*; y*_*r3*_ as above:

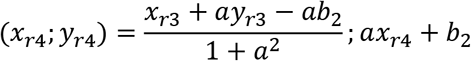

If *a<0*, then *x*_*r1*_*;y*_*r1*_ is set to *x*_*r1*_*;0* (**Fig 3E**). By applying the equation of the line above:

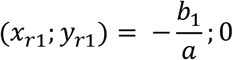

The remaining points of the alignment polygon are subsequently calculated as above.

Finally, to calculate the distance of individual kinetochores away from metaphase, the shortest distance between kinetochores and the metaphase line needs to be computed. To this end, the metaphase line is defined as:

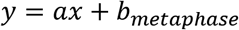

The Midpoint which is the point at which the spindle and metaphase lines intersect is then calculated and used to determine the metaphase line. Thus:

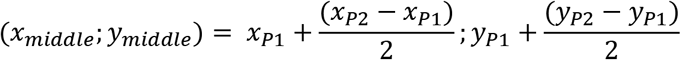

and

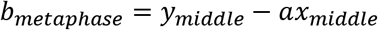

Therefore, the distance *d* of any given point (*x,* y) reflecting a kinetochore position relative to the metaphase line (**Fig 4G**) can be calculated as follows:

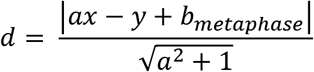

## ACKNOWLEDGEMENTS

We are grateful to all the members of the Elowe lab who have used the methods presented here and have helped optimize it through discussion. This work was supported by Canadian Institutes of Health Research grants to SE (project grant 376557 and GOOP244442). SE holds an FRQS (Fonds de Recherche de Santé Québec) Senior researcher salary award. LGB has been supported by training awards from Desjardins, PROTEO, and “Centre de Recherche sur le Cancer de l’Université Laval”. DKP is currently supported by a CRCHU de Québec training award, and PKS by a PROTEO PhD student award.

## Notes

https://github.com/Elowesab/elowelab

### Competing Interest Statement

The authors have declared no competing interest.

## REFERENCES

Barisic, M., Aguiar, P., Geley, S., and Maiato, H. (2014). Kinetochore motors drive congression of peripheral polar chromosomes by overcoming random arm-ejection forces. Nat Cell Biol 16, 1249–1256.

Bennett, A., Bechi, B., Tighe, A., Thompson, S., Procter, D.J., and Taylor, S.S. (2015). Cenp-E inhibitor GSK923295: Novel synthetic route and use as a tool to generate aneuploidy. Oncotarget 6, 20921–20932.

Cheeseman, I.M. (2014). The kinetochore. Cold Spring Harb Perspect Biol 6, a015826.

Chmatal, L., Yang, K., Schultz, R.M., and Lampson, M.A. (2015). Spatial Regulation of Kinetochore Microtubule Attachments by Destabilization at Spindle Poles in Meiosis I. Curr Biol 25, 1835–1841.

Cimini, D., Wan, X., Hirel, C.B., and Salmon, E.D. (2006). Aurora Kinase Promotes Turnover of Kinetochore Microtubules to Reduce Chromosome Segregation Errors. Current Biology 16, 1711–1718.

Ditchfield, C., Johnson, V.L., Tighe, A., Ellston, R., Haworth, C., Johnson, T., Mortlock, A., Keen, N., and Taylor, S.S. (2003). Aurora B couples chromosome alignment with anaphase by targeting BubR1, Mad2, and Cenp-E to kinetochores. J Cell Biol 161, 267–280.

Espeut, J., and Abrieu, A. (2015). Down-Regulating CENP-E Activity: For Better or for Worse. In Kinesins and Cancer, F.K. Fsb, ed. (Springer Netherlands), pp. 87–99.

Foley, E.A., Maldonado, M., and Kapoor, T.M. (2011). Formation of stable attachments between kinetochores and microtubules depends on the B56-PP2A phosphatase. Nat Cell Biol 13, 1265–1271.

Fonseca, C., and Stumpff, J. (2016). Quantification of Mitotic Chromosome Alignment. Methods Mol Biol 1413, 253–262.

Gordon, D.J., Resio, B., and Pellman, D. (2012). Causes and consequences of aneuploidy in cancer. Nat Rev Genet 13, 189–203.

Harborth, J., Elbashir, S.M., Bechert, K., Tuschl, T., and Weber, K. (2001). Identification of essential genes in cultured mammalian cells using small interfering RNAs. Journal of Cell Science 114, 4557–4565.

Hauf, S., Cole, R.W., LaTerra, S., Zimmer, C., Schnapp, G., Walter, R., Heckel, A., van Meel, J., Rieder, C.L., and Peters, J.-M. (2003). The small molecule Hesperadin reveals a role for Aurora B in correcting kinetochore–microtubule attachment and in maintaining the spindle assembly checkpoint. J Cell Biol 161, 281–294.

Hindriksen, S., Lens, S.M.A., and Hadders, M.A. (2017). The Ins and Outs of Aurora B Inner Centromere Localization. Front Cell Dev Biol 5, 112.

Hinshaw, S.M., and Harrison, S.C. (2018). Kinetochore Function from the Bottom Up. Trends Cell Biol 28, 22–33.

Joglekar, A.P., and Kukreja, A.A. (2017). How Kinetochore Architecture Shapes the Mechanisms of Its Function. Curr Biol 27, R816–R824.

Kapoor, T.M., Lampson, M.A., Hergert, P., Cameron, L., Cimini, D., Salmon, E.D., McEwen, B.F., and Khodjakov, A. (2006). Chromosomes Can Congress to the Metaphase Plate Before Biorientation. Science 311, 388–391.

Kuniyasu, K., Iemura, K., and Tanaka, K. (2018). Delayed Chromosome Alignment to the Spindle Equator Increases the Rate of Chromosome Missegregation in Cancer Cell Lines. Biomolecules 9.

Lampson, M.A., and Grishchuk, E.L. (2017). Mechanisms to Avoid and Correct Erroneous Kinetochore-Microtubule Attachments. Biology 6, 1.

Lampson, M.A., and Kapoor, T.M. (2005). The human mitotic checkpoint protein BubR1 regulates chromosome–spindle attachments. Nat Cell Biol 7, 93–98.

Liu, D., Vleugel, M., Backer, C.B., Hori, T., Fukagawa, T., Cheeseman, I.M., and Lampson, M.A. (2010). Regulated targeting of protein phosphatase 1 to the outer kinetochore by KNL1 opposes Aurora B kinase. J Cell Biol 188, 809–820.

Maiato, H., Gomes, A.M., Sousa, F., and Barisic, M. (2017). Mechanisms of Chromosome Congression during Mitosis. Biology 6, 13.

McGranahan, N., Burrell, R.A., Endesfelder, D., Novelli, M.R., and Swanton, C. (2012). Cancer chromosomal instability: therapeutic and diagnostic challenges. EMBO Rep 13, 528–538.

Meppelink, A., Kabeche, L., Vromans, M.J.M., Compton, D.A., and Lens, S.M.A. (2015). Shugoshin-1 balances Aurora B kinase activity via PP2A to promote chromosome bi-orientation. Cell Rep 11, 508–515.

Musacchio, A., and Desai, A. (2017). A Molecular View of Kinetochore Assembly and Function. Biology (Basel) 6.

Nijenhuis, W., Vallardi, G., Teixeira, A., Kops, G.J.P.L., and Saurin, A.T. (2014). Negative feedback at kinetochores underlies a responsive spindle checkpoint signal. Nat Cell Biol 16, 1257–1264.

Schaar, B.T., Chan, G.K.T., Maddox, P., Salmon, E.D., and Yen, T.J. (1997). CENP-E Function at Kinetochores Is Essential for Chromosome Alignment. J Cell Biol 139, 1373–1382.

Shrestha, Roshan L., and Draviam, Viji M. (2013). Lateral to End-on Conversion of Chromosome-Microtubule Attachment Requires Kinesins CENP-E and MCAK. Current Biology 23, 1514–1526.

Soto, M., Raaijmakers, J.A., and Medema, R.H. (2019). Consequences of Genomic Diversification Induced by Segregation Errors. Trends in Genetics 35, 279–291.

Storchova, Z. (2018). Evolution of aneuploidy: overcoming the original CIN. Genes Dev 32, 1459–1460.

Tanudji, M., Shoemaker, J., L’Italien, L., Russell, L., Chin, G., and Schebye, X.M. (2004). Gene silencing of CENP-E by small interfering RNA in HeLa cells leads to missegregation of chromosomes after a mitotic delay. Mol Biol Cell 15, 3771–3781.

Weaver, B.A.A., Silk, A.D., Montagna, C., Verdier-Pinard, P., and Cleveland, D.W. (2007). Aneuploidy acts both oncogenically and as a tumor suppressor. Cancer Cell 11, 25–36.

Welburn, J.P.I., Vleugel, M., Liu, D., Yates, J.R., Lampson, M.A., Fukagawa, T., and Cheeseman, I.M. (2010). Aurora B Phosphorylates Spatially Distinct Targets to Differentially Regulate the Kinetochore-Microtubule Interface. Molecular Cell 38, 383–392.

Ye, A.A., Deretic, J., Hoel, C.M., Hinman, A.W., Cimini, D., Welburn, J.P., and Maresca, T.J. (2015). Aurora A Kinase Contributes to a Pole-Based Error Correction Pathway. Curr Biol 25, 1842–1851.

Yen, T.J., Compton, D.A., Wise, D., Zinkowski, R.P., Brinkley, B.R., Earnshaw, W.C., and Cleveland, D.W. (1991). CENP-E, a novel human centromere-associated protein required for progression from metaphase to anaphase. EMBO J 10, 1245–1254.

Yu, K.-W., Zhong, N., Xiao, Y., and She, Z.-Y. (2019). Mechanisms of kinesin-7 CENP-E in kinetochore–microtubule capture and chromosome alignment during cell division. Biology of the Cell 111, 143–160.

